# KHNYN is essential for ZAP-mediated restriction of HIV-1 containing clustered CpG dinucleotides

**DOI:** 10.1101/581785

**Authors:** Mattia Ficarelli, Harry Wilson, Rui Pedro Galão, Stuart J D Neil, Chad M Swanson

## Abstract

CpG dinucleotides are suppressed in most vertebrate RNA viruses, including HIV-1, and introducing CpGs into RNA virus genomes inhibits their replication. The zinc-finger antiviral protein (ZAP) binds regions of viral RNA containing CpGs and targets them for degradation. ZAP does not have enzymatic activity and recruits other cellular proteins to inhibit viral replication. Here we show that KHNYN, a protein with no previously known function, interacts with ZAP. KHNYN overexpression selectively inhibits HIV-1 containing clustered CpG dinucleotides and this requires ZAP and its cofactor TRIM25. KHNYN requires both its KH-like domain and NYN endonuclease domain for antiviral activity. Crucially, depletion of KHNYN eliminated the deleterious effect of CpG dinucleotides on HIV-1 RNA abundance and infectious virus production indicating that KHNYN is required for this antiviral pathway. Overall, we have identified KHNYN as a novel ZAP cofactor that is essential for innate immune destruction of CpG containing viral RNA.

## Introduction

A major component of the innate immune system are cell intrinsic antiviral proteins. These act at multiple steps in viral replication cycles and some are induced by type I interferons (Schneider et al., 2014). Many viruses have evolved mechanisms to evade inhibition by these proteins. First, viruses can encode proteins that counteract specific antiviral factors. Second, viral protein sequences can evolve to prevent recognition by these factors. An example of these two mechanisms in HIV-1 are the accessory proteins Vif and Vpu that respectively counteract APOBEC3 cytosine deaminases and Tetherin as well as changes in its capsid protein that allow it to at least partially evade human TRIM5*α* (Malim and Bieniasz, 2012). Third, viral nucleic acid sequences can evolve to eliminate motifs that stimulate the innate immune response. The abundance of CpG dinucleotides is suppressed in many vertebrate RNA virus genomes and when CpGs are experimentally introduced into picornaviruses or influenza A, viral replication is inhibited (Atkinson et al., 2014; Burns et al., 2009; Gaunt et al., 2016; Karlin et al., 1994; Tulloch et al., 2014). This shows that CpG suppression in diverse RNA viruses is required for efficient replication.

CpG dinucleotides are also suppressed in the HIV-1 genome and multiple studies have shown that they are deleterious for replication (Antzin-Anduetza et al., 2017; Kypr et al., 1989; Shpaer and Mullins, 1990; Takata et al., 2017; Theys et al., 2018; Wasson et al., 2017). Recently, the cellular antiviral protein ZAP was shown to bind regions of viral RNA with high CpG abundance and target them for degradation, which at least partly explains why this dinucleotide inhibits viral replication (Takata et al., 2017). ZAP (also known as ZC3HAV1) is a component of the innate immune response targeting viral RNAs in the cytoplasm to prevent viral protein synthesis (Li et al., 2015).

ZAP inhibits the replication of a diverse range of viruses including retroviruses, alphaviruses, filoviruses, hepatitis B virus and Japanese encephalitis virus as well as retroelements (Bick et al., 2003; Chiu et al., 2018; Gao et al., 2002; Goodier et al., 2015; Mao et al., 2013; Moldovan and Moran, 2015; Muller et al., 2007; Takata et al., 2017; Zhu et al., 2011). There are two human ZAP isoforms, ZAP-L and ZAP-S (Kerns et al., 2008). Both isoforms contain a N-terminal RNA binding domain containing four CCCH-type zinc finger motifs but ZAP-L also contains a catalytically inactive C-terminal poly(ADP-ribose) polymerase (PARP)-like domain (Chen et al., 2012; Guo et al., 2004; Kerns et al., 2008). Importantly, neither isoform of ZAP has nuclease activity and it likely recruits other cellular proteins to degrade viral RNAs. Identifying and characterizing these cofactors for ZAP is essential to understand how it restricts viral replication.

ZAP requires the E3 ubiquitin ligase TRIM25 for its antiviral activity and TRIM25 depletion reduced the ability of ZAP to inhibit infectious virus production of HIV-1 with clustered CpGs (Li et al., 2017; Takata et al., 2017; Zheng et al., 2017). While ZAP has been reported to interact with several components of the 5’-3’ and 3’-5’ RNA degradation pathways, depletion of these proteins did not substantially increase infectious virus production for HIV-1 containing clustered CpG dinucleotides (Goodier et al., 2015; Guo et al., 2007; Takata et al., 2017; Zhu et al., 2011). This suggests that additional proteins may be required for ZAP to inhibit viral replication. Herein, we identify KHNYN as a cytoplasmic protein that interacts with ZAP and is necessary for CpG dinucleotides to inhibit HIV-1 RNA and protein abundance.

## Results

### KHNYN interacts with ZAP and selectively inhibits HIV-1 containing clustered CpG dinucleotides in a ZAP- and TRIM25-dependent manner

To identify candidate interaction partners for ZAP, a yeast two-hybrid screen was performed for full-length ZAP-S and ZAP-L using prey fragments from an interferon and TLR-ligand induced human macrophage cDNA library. Candidate interacting proteins were assigned a Predicted Biological Score (PBS) from A to F: A = very high confidence in the interaction, B = high confidence in the interaction and C = good confidence in the interaction. Scores of D to F are moderate to low confidence interactions. For ZAP-S, 10 clones containing a prey fragment encoding KHNYN (Figure 1) were isolated and these had a PBS = A. For ZAP-L, two positive clones were identified encoding KHNYN and had a PBS = C. KHNYN has two isoforms (KHNYN-1 and KHNYN-2) that contain a N-terminal KH-like domain and a C-terminal NYN endoribonuclease domain (Figure 1) (Anantharaman and Aravind, 2006). The selected interaction domain, which is the amino acid sequence shared by all prey fragments matching KHNYN, comprised amino acids 572-719 of KHNYN-2 for the clones identified in both screens. A yeast two-hybrid screen was then performed using the same library with full length KHNYN-2 as the bait. Nine clones were isolated that encode ZAP and these had a PBS = A. The selected interaction domain was amino acids 4-352, which is present in both isoforms (Figure 1). Supporting the reproducibility of this interaction, KHNYN has also been identified as a ZAP-interacting factor in large-scale affinity purification–mass spectrometry and *in vivo* proximity-dependent biotinylation (BioID) screens (Huttlin et al., 2017; Youn et al., 2018).

**Figure 1.**
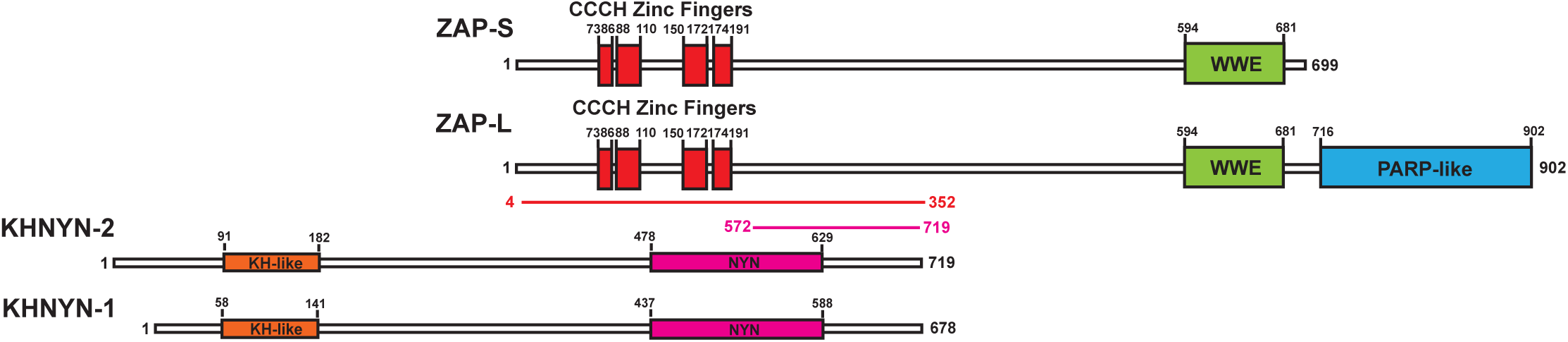
KHNYN is a ZAP-interacting factor identified by yeast two-hybrid screening. A yeast two-hybrid screen for ZAP-S and ZAP-L interacting factors identified a region in KHNYN-1 and KHNYN-2. The selected interaction domain (SID) is the amino acid sequence shared by all prey fragments and is shown in magenta. A reciprocal yeast two-hybrid screen using KHNYN-2 as the bait identified a region in ZAP-S and ZAP-L. The SID is shown in red.

We first confirmed the interaction between ZAP and KHNYN by co-immunoprecipitation and found both KHNYN isoforms interacted with both isoforms of ZAP (Figure 2A and 2B). This interaction was RNase insensitive (Figure 2C). Since ZAP mediates degradation of HIV-1 RNAs with clustered CpG dinucleotides in the cytoplasm (Takata et al., 2017), its cofactors are likely to be localized in this compartment. Therefore, we analyzed the subcellular localization of KHNYN and observed that it localizes to the cytoplasm similar to ZAP (Figure 2D). Its localization was not affected when ZAP was knocked out using CRISPR-Cas9-mediated genome editing.

**Figure 2.**
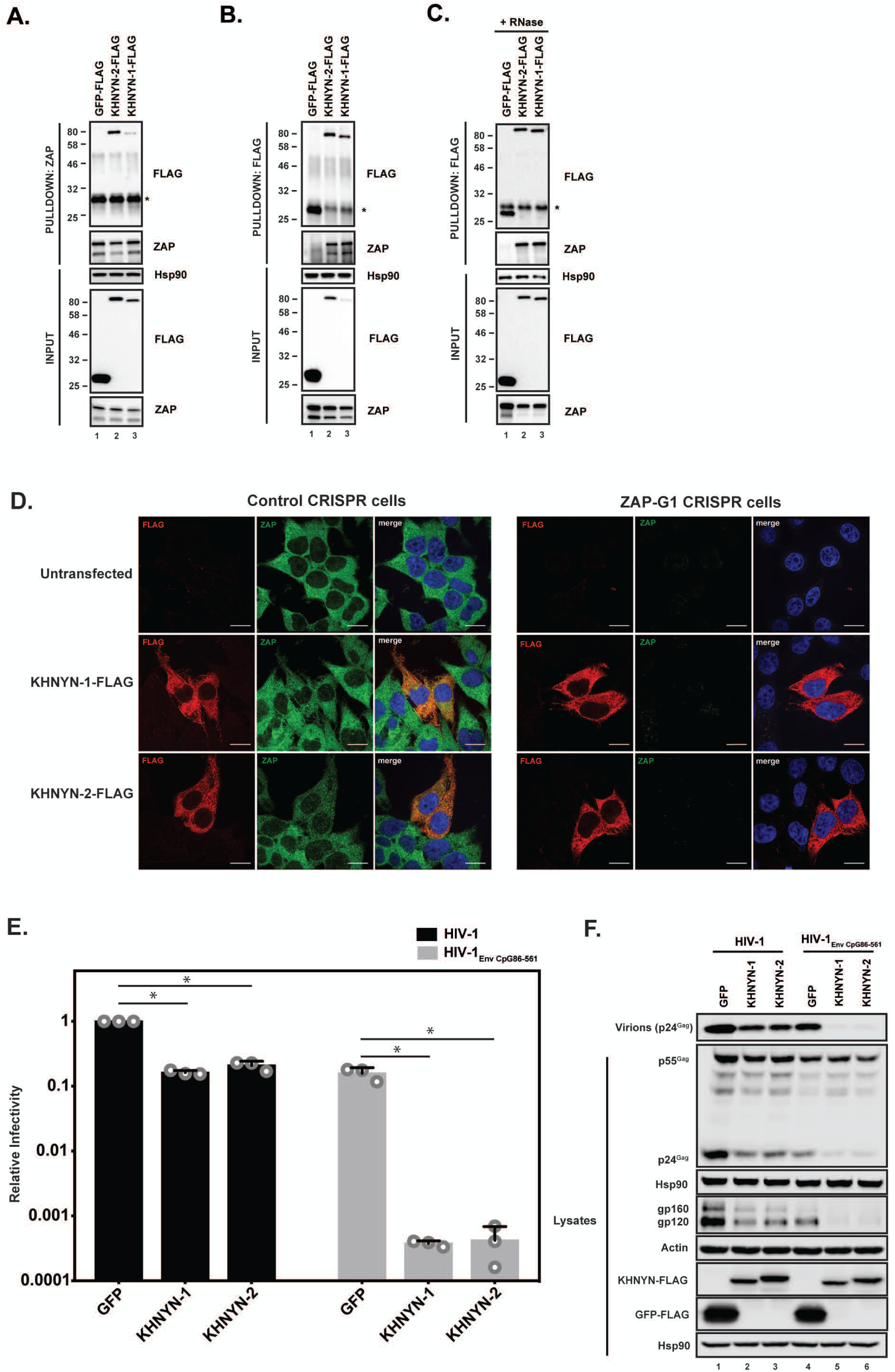
KHNYN interacts with ZAP and selectively inhibits HIV-1 containing clustered CpG dinucleotides. **(A)** Lysates of HEK293T cells transfected either with pGFP-FLAG, pKHNYN-1-FLAG or pKHNYN-2-FLAG were immunoprecipitated with an anti-ZAP antibody. Post-nuclear supernatants and immunoprecipitation samples were analyzed by immunoblotting for Hsp90, KHNYN-FLAG and ZAP. * indicates a non-specific band. **(B)** Lysates of HEK293T cells transfected either with pGFP-FLAG, pKHNYN-1-FLAG or pKHNYN-2-FLAG were immunoprecipitated with an anti-FLAG antibody. Post-nuclear supernatants and immunoprecipitation samples were analyzed by immunoblotting for HSP90, KHNYN-FLAG and ZAP. * indicates a non-specific band. **(C)** Lysates of HEK293T cells transfected with pZAP-L and either pGFP-FLAG, pKHNYN-1-FLAG or pKHNYN-2-FLAG were treated with RNase and then immunoprecipitated with an anti-FLAG antibody. Post-nuclear supernatants and immunoprecipitation samples were analyzed by immunoblotting for HSP90, KHNYN-FLAG and ZAP. * indicates a non-specific band. **(D)** Panels show representative fields for the localization of KHNYN-1-FLAG or KHNYN-2-FLAG and endogenous ZAP in either 293T Control CRISPR cells expressing a guide RNA targeting *LacZ* or 293T *ZAP* guide 1 (ZAP-G1) CRISPR cells. Cells were stained with an anti-FLAG antibody (red), anti-ZAP antibody (green) and DAPI (blue). The scale bar represents 10 µM. **(E-F)** HeLa cells were transfected with 500 ng pHIV-1 or pHIV-1_EnvCpG86-561_ and 500 ng of pGFP-FLAG, pKHNYN-1-FLAG or pKHNYN-2-FLAG. See also Figure 2–figure supplement 1. Culture supernatants were used to infect TZM-bl reporter cells to measure infectivity **(E)**. The bar charts show the average values of three independent experiments normalized to the value obtained for HeLa cells co-transfected with pHIV-1 and pGFP-FLAG. Data are represented as mean ± SD. *p < 0.05 as determined by a two-tailed unpaired t-test. p-values for GFP verses KHNYN-1 and KHNYN-2 for wild type HIV-1 are 2.76 x 10^-9^ and 2.20 x 10^-6^ respectively. p-values for GFP verses KHNYN-1 and KHNYN-2 for HIV-1_EnvCpG86-561_ are 1.50 x 10^-3^ and 1.51 x 10^-3^, respectively. Gag expression in the media as well as Gag, Hsp90, Env, Actin, KHNYN-FLAG and GFP-FLAG expression in the cell lysates was detected using quantitative immunoblotting **(F)**.

The mechanisms that allow a virus to escape the innate immune response often have to be mutated to study the effect of antiviral proteins. For example, HIV-1 Vpu or Vif have be mutated to allow tetherin or APOBEC3 antiviral activity to be analyzed (Malim and Bieniasz, 2012). Since CpG dinucleotides are suppressed in HIV-1, endogenous ZAP does not target the wild type virus (Takata et al., 2017). However, a ZAP-sensitive HIV-1 can be created by introducing CpGs through synonymous mutations into the *env* region of the viral genome. This makes HIV-1 an excellent system to study the mechanism of action of this antiviral protein because isogenic viruses can be analyzed that differ only in their CpG abundance and therefore ZAP-sensitivity (Takata et al., 2017). To determine if KHNYN overexpression inhibited wild type HIV-1 or HIV-1 with 36 CpG dinucleotides introduced into *env* nucleotides 86-561 (HIV-1_EnvCpG86-561_) (Figure 2–figure supplement 1), each isoform was overexpressed in the context of a single cycle infectivity assay. As expected, transfection of the HIV-1_EnvCpG86-561_ provirus into HeLa cells yielded substantially less infectious virus than wild type HIV-1, which was accounted for by reduced expression of Gag and Env proteins (Figure 2E and 2F). While KHNYN-1 or KHNYN-2 overexpression decreased wild type HIV-1 infectivity by ∼5-fold, they decreased HIV-1_EnvCpG86-561_ infectivity by ∼400-fold (Figure 2E). The inhibition of infectivity by KHNYN-1 or KHNYN-2 correlated with further decreases in Gag expression, Env expression, and virion production (Figure 2F). Overall, KHNYN appeared to selectively inhibit HIV-1_EnvCpG86-561_ infectious virion production.

We then determined whether ZAP is necessary for KHNYN to inhibit HIV-1 with clustered CpG dinucleotides. Control or ZAP knockout cells (Figure 3A) were co-transfected with pHIV-1 or pHIV-1_EnvCpG86-561_ and increasing amounts of pKHNYN-1. Wild type HIV-1 infectious virus production was not affected by ZAP depletion and HIV-1_EnvCpG86-561_ infectivity was restored in ZAP knockout cells (Figure 3B, 0 ng of KHNYN-1), confirming that ZAP is necessary to inhibit HIV-1 with CpGs into *env* (Takata et al., 2017). At low levels of KHNYN-1 overexpression (such as 62.5 ng), there was no substantial decrease in infectivity for wild type HIV-1 while HIV-1_EnvCpG86-561_ infectivity was inhibited in a ZAP-dependent manner (Figure 3B and 4A). The decrease in infectivity for HIV-1_EnvCpG86-561_ in control cells transfected with pKHNYN-1 correlated with decreases in Gag expression, Env expression and virion production (Figure 3C).

**Figure 3.**
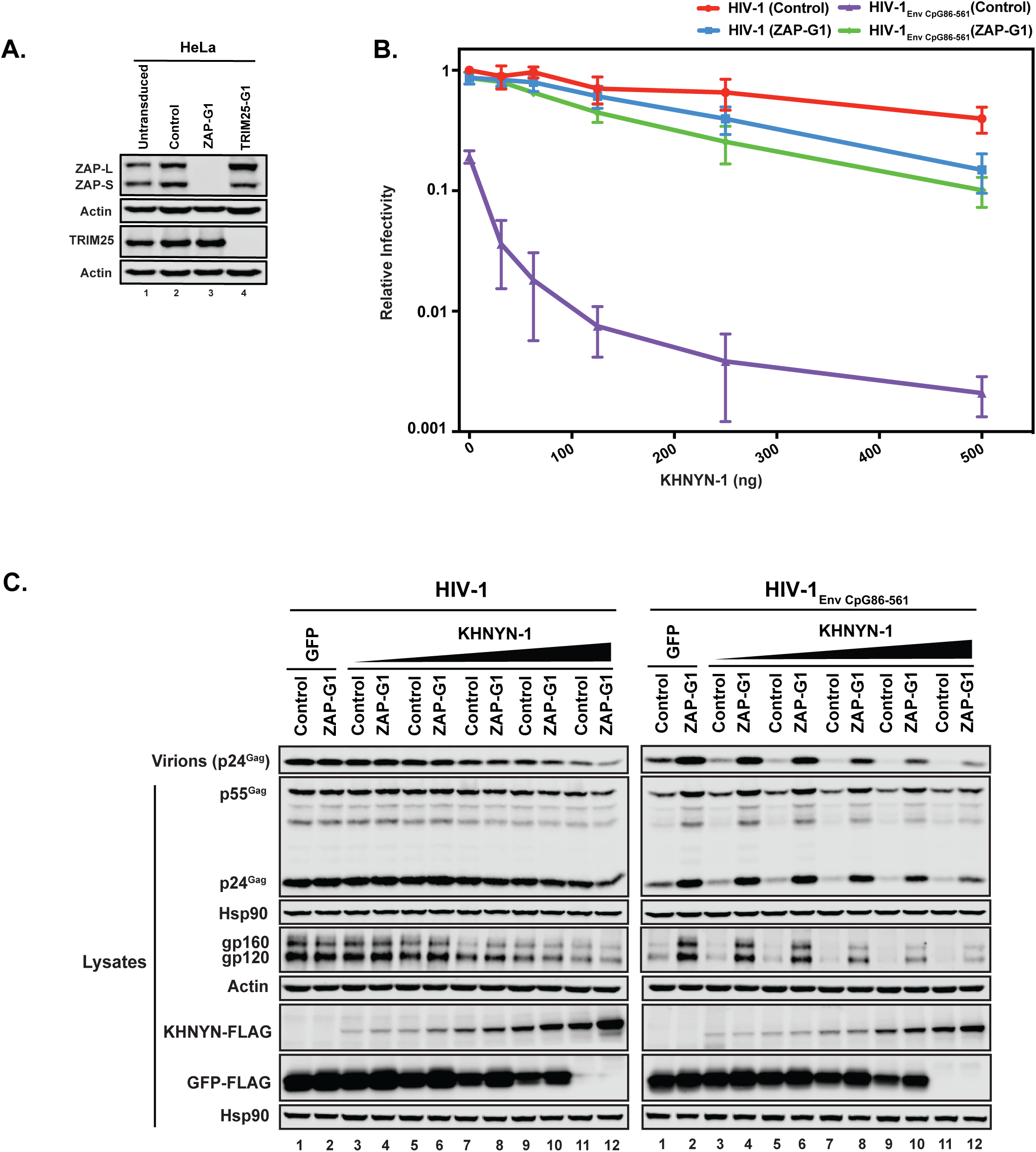
ZAP is required for KHNYN to inhibit infectious virion production for HIV-1 with clustered CpG dinucleotides. **(A)** ZAP, TRIM25 and Actin expression in HeLa cells, HeLa Control CRISPR cells (expressing a guide RNA targeting the *firefly luciferase* gene), HeLa *ZAP* CRISPR guide 1 (ZAP-G1) cells and HeLa *TRIM25* CRISPR guide 1 (TRIM25-G1) cells were detected using quantitative immunoblotting. **(B-C)** HeLa Control CRISPR cells or ZAP-G1 CRISPR cells were transfected with 500 ng pHIV-1 or pHIV-1_EnvCpG86-561_ and 500 ng of pGFP-FLAG or 31.25 ng, 62.5 ng, 125 ng, 250 ng or 500 ng pKHNYN-1-FLAG plus the amount of pGFP-FLAG required to make 500 ng total. Culture supernatants from the cells were used to infect TZM-bl reporter cells **(B)**. Each point shows the average value of three independent experiments normalized to the value obtained for HeLa Control CRISPR cells co-transfected with pHIV-1 and pGFP-FLAG. Data are represented as mean ± SD. Gag expression in the media as well as Gag, Hsp90, Env, Actin, KHNYN-FLAG and GFP-FLAG expression in the cell lysates was detected using quantitative immunoblotting **(C)**.

Next, we analyzed how ZAP and KHNYN regulate HIV-1 genomic RNA abundance in the cell lysate and media. As expected (Takata et al., 2017), HIV-1_EnvCpG86-561_ genomic RNA abundance was decreased in control cells but was similar to wild type HIV-1 in ZAP knockout cells (Figure 4B-C, compare GFP samples). In control cells, 62.5 ng of KHNYN-1 or KHNYN-2 inhibited HIV-1_EnvCpG86-561_ genomic RNA abundance compared to the GFP control (Figure 4B-C). Importantly, KHNYN-1 and KHNYN-2 did not affect wild type HIV-1 genomic RNA levels and did not substantially inhibit HIV-1_EnvCpG86-561_ genomic RNA abundance in ZAP knockout cells. This demonstrates that KHNYN targets HIV-1 RNA containing clustered CpG dinucleotides in a ZAP-dependent manner.

**Figure 4.**
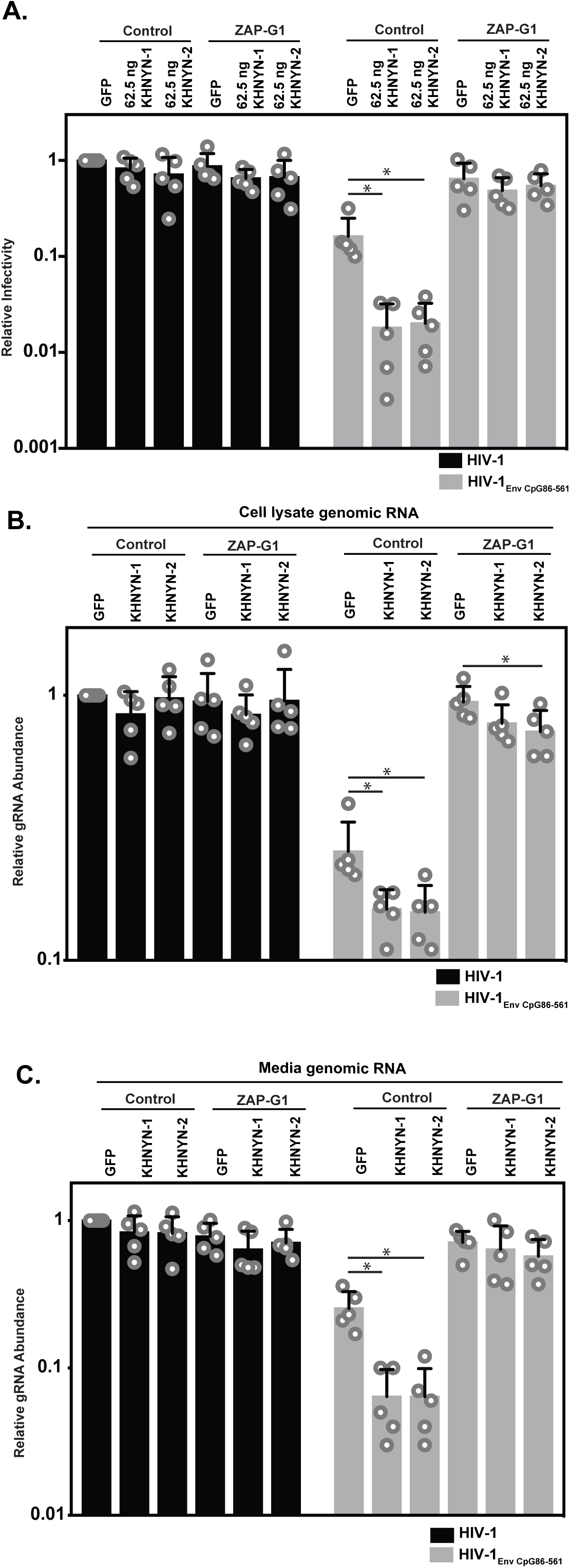
ZAP is required for KHNYN to inhibit genomic RNA abundance for HIV-1 with clustered CpG dinucleotides. **(A-C)** HeLa Control CRISPR cells or ZAP-G1 CRISPR cells were transfected with 500 ng pHIV-1 or pHIV-1_EnvCpG86-561_ and 500 ng of pGFP-FLAG or 62.5 ng pKHNYN-1/2-FLAG plus 437.5 ng of pGFP-FLAG. Culture supernatants from the cells were used to infect TZM-bl reporter cells **(A)**. The bar chart shows the average value of five independent experiments normalized to the value obtained for HeLa Control CRISPR cells co-transfected with pHIV-1 and pGFP-GFP. Data are represented as mean ± SD. *p < 0.05 as determined by a two-tailed unpaired t-test. p-values for GFP verses KHNYN-1 and KHNYN-2 for HIV-1_EnvCpG86-561_ in Control cells are 6.78 x 10^-3^ and 7.20 x 10^-3^, respectively. p-values for GFP verses KHNYN-1 and KHNYN-2 for HIV-1_EnvCpG86-561_ in ZAP-G1 cells are 3.22 x 10^-1^ and 5.33 x 10^-1^, respectively. RNA was extracted from cell lysates **(B)** and media **(C)** and genomic RNA (gRNA) abundance was quantified by qRT-PCR. The bar charts show the average value of five independent experiments normalized to the value obtained for HeLa Control CRISPR cells co-transfected with pHIV-1 and pGFP-GFP. Data are represented as mean ± SD. *p < 0.05 as determined by a two-tailed unpaired t-test. For HIV-1_EnvCpG86-561_ genomic RNA in Control cell lysates, the GFP verses KHNYN-1 and KHNYN-2 p-values are 2.14 x 10^-2^ and 2.30 x 10^-2^, respectively. For HIV-1_EnvCpG86-561_ genomic RNA in ZAP- G1 cell lysates, the GFP verses KHNYN-1 and KHNYN-2 p-values are 1.01 x 10^-1^ and 4.33 x 10^-2^, respectively. For HIV-1_EnvCpG86-561_ genomic RNA in Control cell media, p- values for GFP verses KHNYN-1 and KHNYN-2 are 8.97 x 10^-4^ and 9.38 x 10^-4^, respectively. For HIV-1_EnvCpG86-561_ genomic RNA in ZAP-G1 cell media, p-values for GFP verses KHNYN-1 and KHNYN-2 are 6.09 x 10^-1^ and 1.87 x 10^-1^ respectively.

TRIM25 is required for ZAP to bind viral RNA and its antiviral activity, though the mechanism by which it regulates ZAP is unclear (Li et al., 2017; Zheng et al., 2017). To determine if TRIM25 is necessary for the antiviral activity of KHNYN, 62.5 ng of pKHNYN-1 or pKHNYN-2 was co-transfected with pHIV-1 or pHIV-1_EnvCpG86-561_ in control and TRIM25 knockout cells. Both isoforms of KHNYN inhibited HIV-1_EnvCpG86-561_ much less potently in TRIM25 knockout cells than control cells and had no effect on wild type HIV-1 in either cell line (Figure 5A-B). One possible reason that TRIM25 is necessary for KHNYN antiviral activity could be that it regulates the interaction between ZAP and KHNYN. We pulled down KHNYN-FLAG and western blotted for ZAP in control and TRIM25 knockdown cells (Figure 5C). Both isoforms of KHNYN pulled down ZAP in both cell lines, indicating that TRIM25 is not required for the interaction between these proteins. Interestingly, KHNYN also pulled down TRIM25 in control cells (Figure 5C). Therefore, we analyzed whether KHNYN interacted with TRIM25 in control and ZAP knockout cells and observed that both isoforms of KHNYN-FLAG immunoprecipitated TRIM25 in the presence and absence of ZAP (Figure 5D). In sum, KHNYN requires TRIM25 to inhibit HIV-1 containing clustered CpG dinucleotides, but TRIM25 is not necessary for the interaction between ZAP and KHNYN. Furthermore, ZAP, KHNYN and TRIM25 appear to be in a complex together.

**Figure 5.**
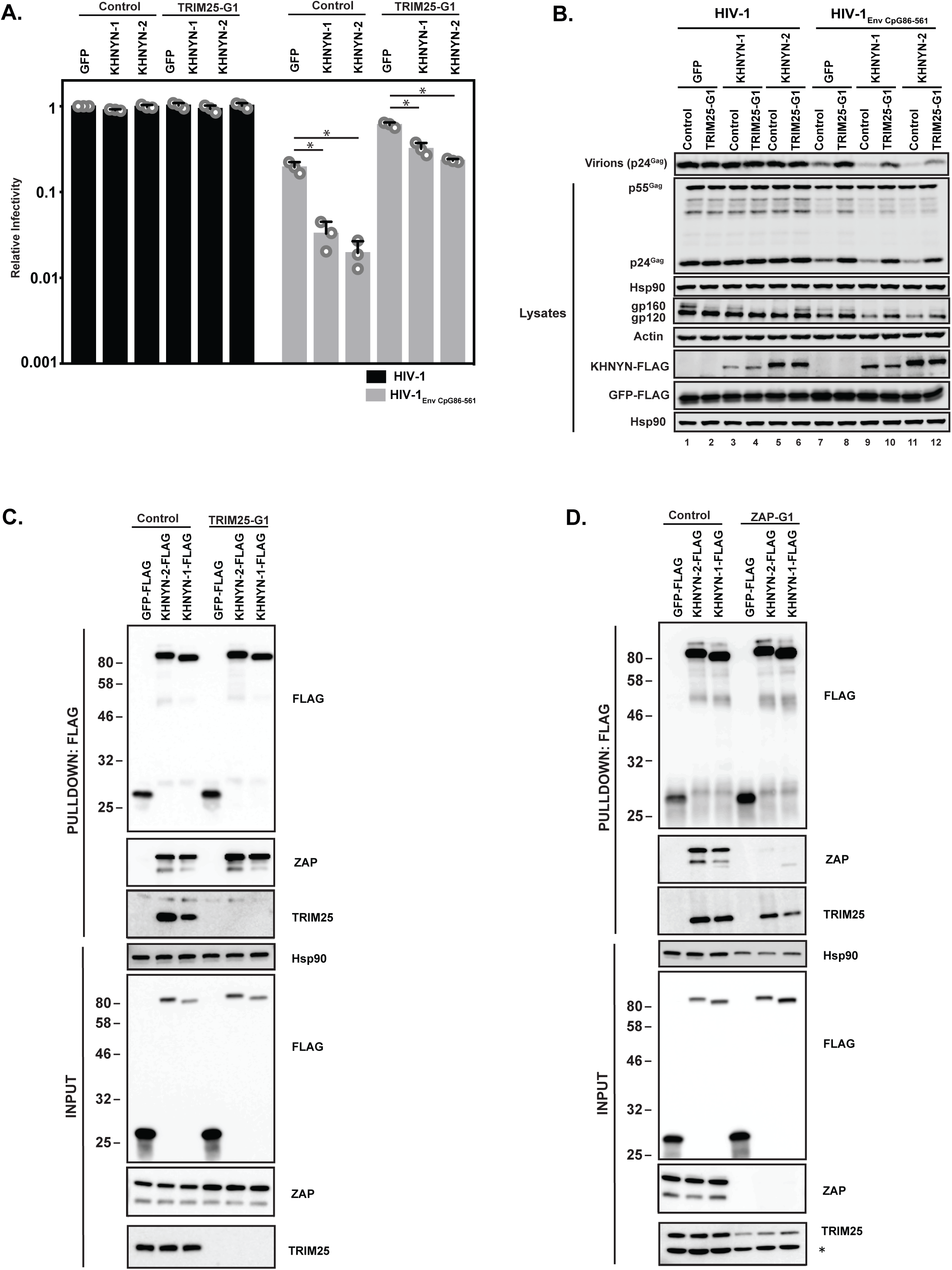
TRIM25 is required for KHNYN to inhibit HIV-1 with clustered CpG dinucleotides. **(A-B)** HeLa Control CRISPR cells or TRIM25-G1 CRISPR cells were transfected with 500 ng pHIV-1 or pHIV-1_EnvCpG86-561_ and 500 ng of pGFP-FLAG or 62.5 ng pKHNYN-1-FLAG or pKHNYN-2-FLAG plus 437.5 ng of pGFP-FLAG. Culture supernatants from the cells were used to infect TZM-bl reporter cells **(A)**. The bar charts show the average value of three independent experiments normalized to the value obtained for HeLa Control CRISPR cells co-transfected with pHIV-1 and pGFP-FLAG. Data are represented as mean ± SD. *p < 0.05 as determined by a two-tailed unpaired t-test. p-values for GFP verses KHNYN-1 and KHNYN-2 for HIV-1_EnvCpG86-561_ in Control cells are 8.95 x 10^-4^ and 5.42 x 10^-4^, respectively. p-values for GFP verses KHNYN-1 and KHNYN-2 for HIV-1_EnvCpG86-561_ in TRIM25-G1 CRISPR cells are 1.78 x 10^-3^ and 1.01 x 10^-4^, respectively. Gag expression in the media as well as Gag, Hsp90, Env, Actin, KHNYN-FLAG and GFP-FLAG expression in the cell lysates was detected using quantitative immunoblotting **(B)**. **(C)** Lysates of Control and TRIM25 CRISPR HEK293T cells transfected with pGFP-FLAG, pKHNYN-1-FLAG or pKHNYN-2-FLAG were immunoprecipitated with an anti-FLAG antibody. Post-nuclear supernatants and immunoprecipitation samples were analyzed by immunoblotting for HSP90, KHNYN-FLAG, TRIM25 and ZAP. The blots are representative of two independent experiments. **(D)** Lysates of Control and ZAP CRISPR HEK293T cells transfected with pGFP-FLAG, pKHNYN-1-FLAG or pKHNYN-2-FLAG were immunoprecipitated with an anti-FLAG antibody. Post-nuclear supernatants and immunoprecipitation samples were analyzed by immunoblotting for HSP90, KHNYN-FLAG, TRIM25 and ZAP. * indicates a non-specific band. The blots are representative of two independent experiments.

### The KH-like and NYN domains are necessary for KHNYN antiviral activity

As its name implies, KHNYN contains a KH-like domain and a NYN domain (Figure 6A). The KH-like domain differs from canonical KH domains due to a potential small metal chelating module containing two cysteines and a histidine inserted into the central region of the domain (Anantharaman and Aravind, 2006). Since this has diverged substantially from a standard KH domain, it has also been called a CGIN1 domain and is only known to be present in two other proteins (Marco and Marin, 2009). While most KH domains bind nucleic acids (Nicastro et al., 2015), the insertion in the KH-like domain in KHNYN may disrupt RNA binding and indicate that it has a different function. To analyze the functional importance of the KH-like domain, we deleted it and found that KHNYN-1ΔKH and KHNYN-2ΔKH had reduced antiviral activity compared to the wild type protein (Figure 6B-C). These mutant proteins localized to the cytoplasm and formed foci that were not present for wild type KHNYN-1 or KHNYN-2 (Figure 6–figure supplement 1).

**Figure 6.**
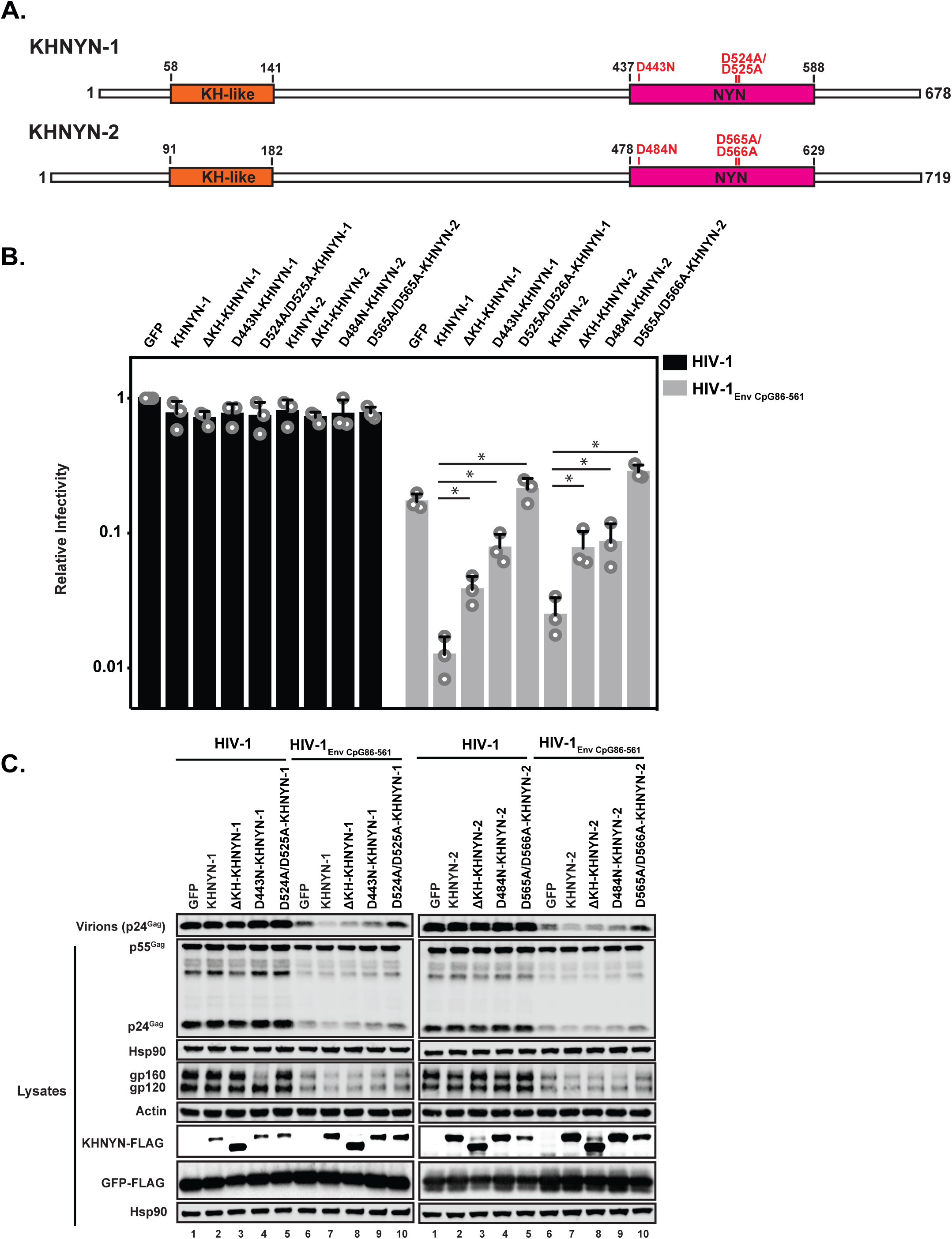
The KH and NYN domains are necessary for KHNYN antiviral activity. **(A)** Schematic of KHNYN-1 and KHNYN-2 domains and mutations in the NYN domain. **(B-C)** HeLa cells were transfected with 500 ng pHIV-1 or pHIV-1_EnvCpG86-561_ and either 500 ng of pGFP-FLAG or 62.5 ng pKHNYN-1/2-FLAG and 437.5 ng of pGFP-FLAG. Culture supernatants were used to infect TZM-bl reporter cells to measure infectivity **(B)**. The bar charts show the average values of three independent experiments normalized to the value obtained for HeLa cells co-transfected with pHIV-1 and pGFP-FLAG. Data are represented as mean ± SD. *p < 0.05 as determined by a two-tailed unpaired t-test. p-values for KHNYN-1 versus ΔKH-KHNYN-1, D443N-KHNYN-1, and D524A/D525A-KHNYN-1 for HIV-1_EnvCpG86-561_ are 1.31 x 10^-2^, 5.22 x 10^-3^, and 1.26 x 10^-3^, respectively. p-values for KHNYN-2 versus ΔKH-KHNYN-2, D443N-KHNYN-2, and D524A/D525A-KHNYN-2 for HIV-1_EnvCpG86-561_ are 2.88 x 10^-2^, 3.26 x 10^-2^, and 2.56 x 10^-4^, respectively. Gag expression in the media as well as Gag, Hsp90, Env, Actin, KHNYN-1/2-FLAG and GFP-FLAG expression in the cell lysates was detected using quantitative immunoblotting **(C)**. See also Figure 6–figure supplement 1 and 2.

NYN domains have endonuclease activity and belong to the PIN nuclease domain superfamily. There are at least seven human proteins with a potentially active NYN domain and they have been structurally characterized in ZC3H12A and MARF1 (Matsushita et al., 2009; Nishimura et al., 2018; Xu et al., 2012; Yao et al., 2018). These domains contain a negatively charged active site with four aspartic acid residues coordinating a magnesium ion, which activates a water molecule for nucleophilic attack of the phosphodiester group on the target RNA. Mutation of these acidic residues inhibits nuclease activity by disrupting the bonds that directly or indirectly interact with the magnesium ion. ZC3H12A (also known as MCPIP1 and Regnase) is a RNA binding protein that, similar to ZAP, contains a CCCH zinc finger domain and degrades cellular and viral RNAs (Takeuchi, 2018). The NYN domain in ZC3H12A has 56% identity to the NYN domain in KHNYN (Figure 6–figure supplement 2A). ZC3H12A containing a D141N mutation in the NYN domain had decreased endonuclease activity and did not degrade RNA containing the IL-6 3’ UTR (Matsushita et al., 2009). MARF1 is required for meiosis and retrotransposon silencing in oocytes and a D426A/D427A mutation inhibited its endoribonuclease activity (Nishimura et al., 2018; Su et al., 2012). We made these equivalent mutations in KHNYN (Figure 6A and Figure 6–figure supplement 2B) and tested their ability to inhibit HIV-1_EnvCpG86-561_ gene expression and infectious virus production (Figure 6B-C). KHNYN-1 D443N and KHNYN-2 D484N had substantially decreased activity against HIV-1_EnvCpG86-561_. Strikingly, KHNYN-1 D524A/D525A and KHNYN-2 D565A/D566A had no antiviral activity even though they were expressed at similar levels to the wild type KHNYN. KHNYN proteins with mutations in the NYN domain localized to the cytoplasm, indicating that disruption of this domain’s activity did not substantially affect their subcellular localization (Figure 6–figure supplement 1).

### KHNYN is necessary for CpG dinucleotides to inhibit HIV-1 RNA and protein expression

To determine if KHNYN is required for CpG dinucleotides to inhibit HIV-1 infectious virus production, we depleted it using CRISPR-Cas9-mediated genome editing. We were unable to identify an antibody that detected endogenous KHNYN. However, FLAG-tagged KHNYN-1 and KHNYN-2 were depleted in bulk population of cells expressing two different CRISPR guide sequences as well as two clonal cell lines from each bulk population (Figure 7A and 7D). Importantly, when the CRISPR PAM sequence was mutated in the KHNYN plasmids, KHNYN-1 and KHNYN-2 were no longer depleted. These KHNYN CRISPR cells were then transfected with pHIV-1 or pHIV-1_EnvCpG86-561_. In the KHNYN CRISPR cells, the CpG dinucleotides in HIV-1_EnvCpG86-561_ no longer inhibited infectious virus production, Gag expression or Env expression (Figure 7B, 7C, 7E and 7F). Wild type HIV-1 Gag expression, Env expression and infectious virus production was not altered in the KHNYN CRISPR cells compared to the control cells.

**Figure 7.**
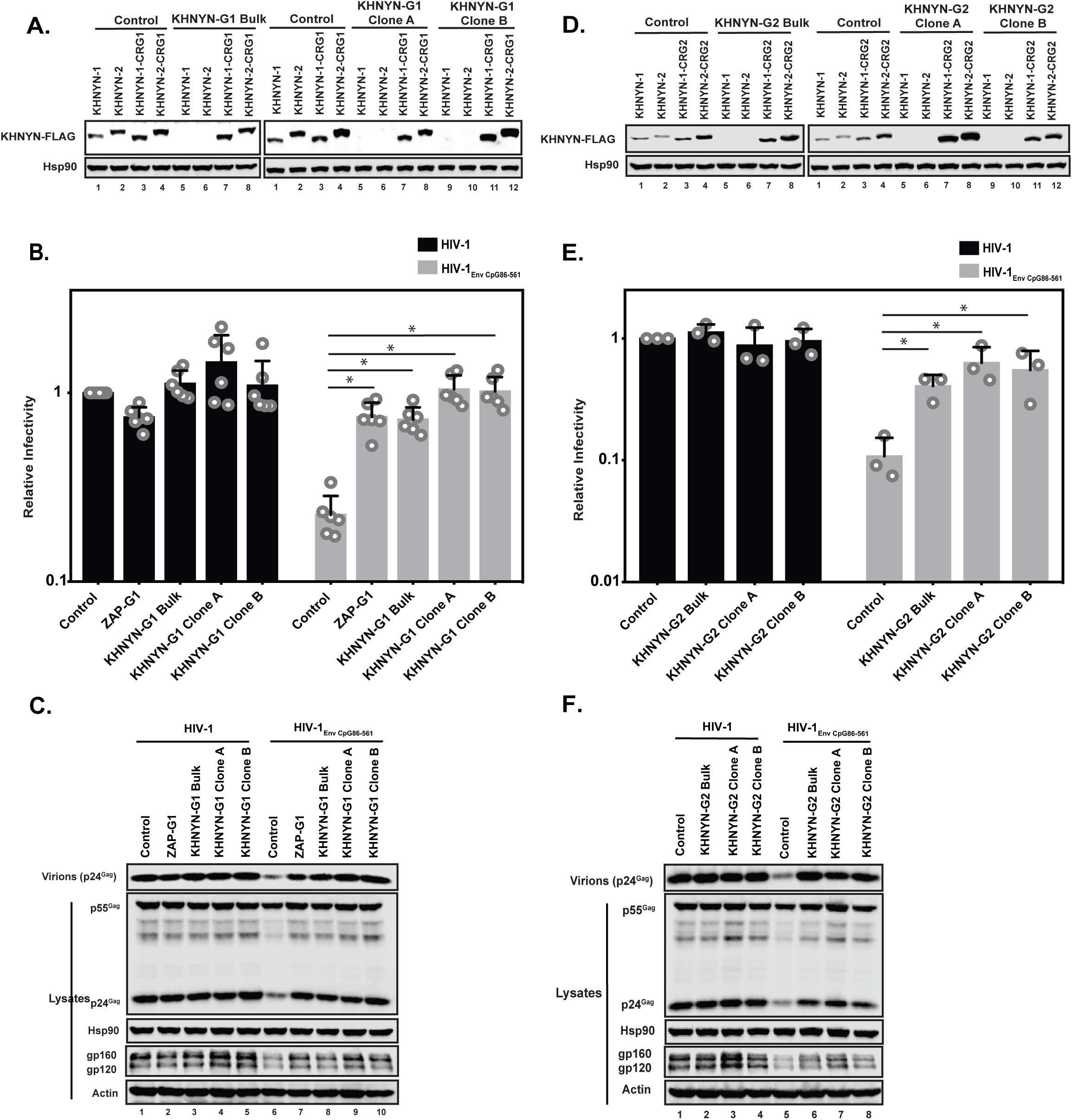
KHNYN is necessary for CpG dinucleotides to inhibit HIV-1 infectious virus production. **(A)** 100 ng pKHNYN-1-FLAG, pKHNYN-2-FLAG, pKHNYN-1-FLAG with a mutation in the PAM that prevents it from being targeted by *KHNYN* guide 1 (pKHNYN-1-CRG1-FLAG) or pKHNYN-2-CRG1-FLAG plus 400 ng of pGFP were transfected into HeLa Control CRISPR cells, *KHNYN* guide 1 (KHNYN-G1) CRISPR bulk cells or KHNYN-G1 CRISPR single cell clones A and B. KHNYN-FLAG abundance was measured by quantitative western blotting. **(B-C)** HeLa Control CRISPR cells, ZAP-G1 CRISPR cells, *KHNYN* guide 1 (KHNYN-G1) CRISPR bulk cell population or single cell clones A and B were transfected with 500 ng pHIV-1 or pHIV-1_EnvCpG86-561_ and 500 ng pGFP. Culture supernatants were used to infect TZM-bl reporter cells to measure infectivity **(B)**. The bar charts show the average value of six independent experiments normalized to the value obtained for HeLa Control CRISPR cells co-transfected with pHIV-1 and pGFP. Data are represented as mean ± SD. *p < 0.05 as determined by a two-tailed unpaired t-test. p-values for HIV-1_EnvCpG86-561_ in Control cells versus ZAP-G1, KHNYN-G1 Bulk, KHNYN-G1 Clone A, and KHNYN-G1 Clone B CRISPR cells are 8.23 x 10^-6^, 2.81 x 10^-6^, 1.60 x 10^-6^, and 2.24 x 10^-6^, respectively. Gag expression in the media as well as Gag, Hsp90, Env, and Actin expression in the cell lysates was detected using quantitative immunoblotting **(C)**. **(D)** 100 ng pKHNYN-1-FLAG, pKHNYN-2-FLAG, pKHNYN-1-FLAG with a mutation in the PAM that prevents it from being targeted by *KHNYN* guide 2 (pKHNYN-1-CRG2-FLAG) or pKHNYN-2-CRG2-FLAG plus 400 ng pGFP were transfected into HeLa Control CRISPR cells, *KHNYN* guide 2 (KHNYN-G2) CRISPR bulk cells or KHNYN-G2 CRISPR single cell clones A and B. KHNYN-FLAG abundance was measured by quantitative immunoblotting. **(E-F)** HeLa Control CRISPR cells, KHNYN-G2 CRISPR bulk cells or KHNYN-G2 CRISPR single cell clones A and B were transfected with 500 ng pHIV-1 or pHIV-1_EnvCpG86-561_ and 500 ng pGFP. Culture supernatants were used to infect TZM-bl reporter cells to measure infectivity **(E)**. The bar charts show the average value of three independent experiments normalized to the value obtained for HeLa Control CRISPR cells co-transfected with pHIV-1 and pGFP. Data are represented as mean ± SD. *p < 0.05 as determined by two-tailed unpaired t-test. p-values for HIV-1_EnvCpG86-561_ in Control cells versus KHNYN-G2 Bulk, KHNYN-G2 Clone A, and KHNYN-G2 Clone B CRISPR cells are 7.98 x 10^-3^, 1.49 x 10^-2^, and 3.44 x 10^-2^, respectively. Gag expression in the media as well as Gag, Hsp90, Env, and Actin expression in the cell lysates was detected using quantitative immunoblotting **(F)**.

We then analyzed HIV-1 genomic RNA abundance in the KHNYN CRISPR cells. Similar to HIV-1 protein expression and infectivity, HIV-1_EnvCpG86-561_ genomic RNA abundance was similar in the in the cell lysate and media to wild type HIV-1 (Figure 8A-B), indicating that the CpG dinucleotides no longer inhibited RNA abundance. The wild type HIV-1 genomic RNA abundance was not altered in the KHNYN CRISPR cells compared to the control cells, further showing the specific effect of KHNYN for viral RNA containing CpG dinucleotides.

**Figure 8.**
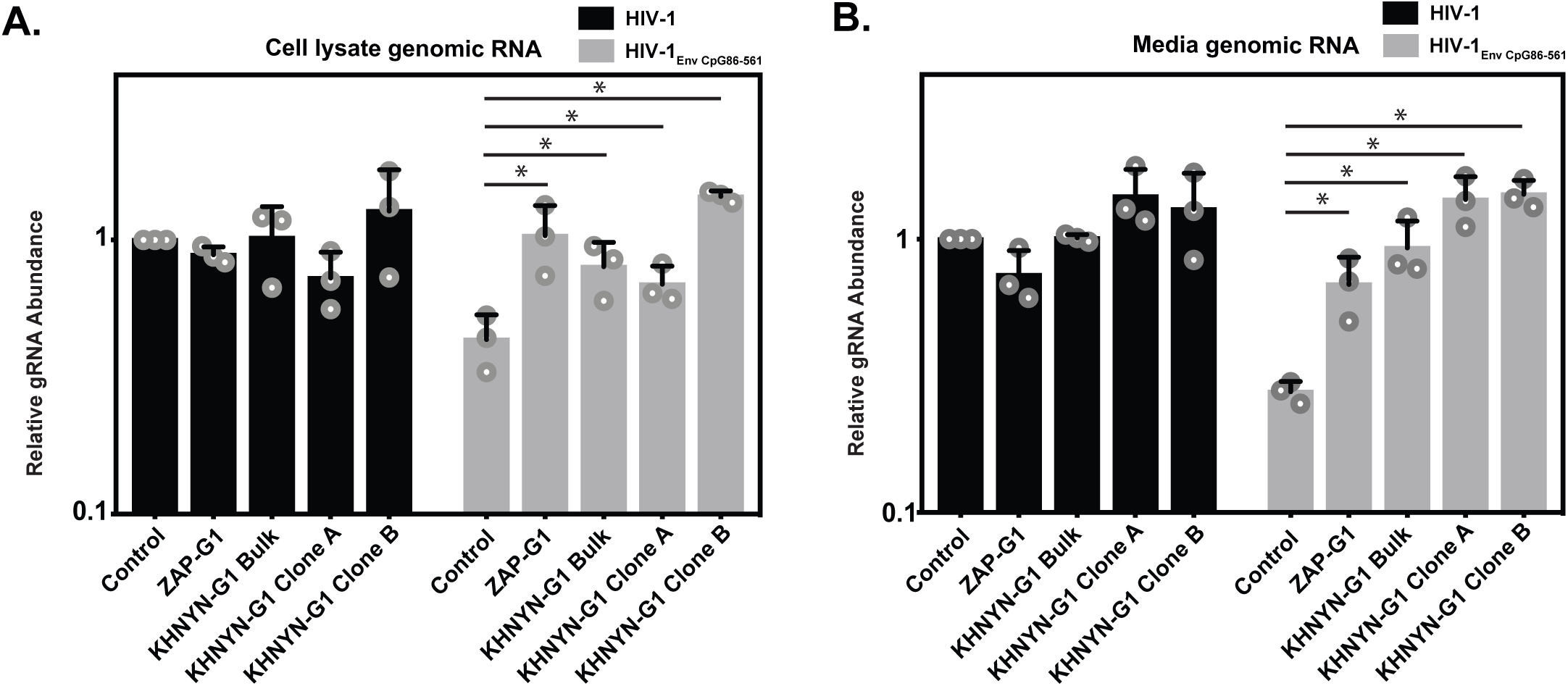
KHNYN is necessary for CpG dinucleotides to inhibit HIV-1 genomic RNA expression. **(A-B)** HeLa Control CRISPR cells, ZAP-G1 CRISPR cells, KHNYN-G1 CRISPR bulk cells or 2 independent clones were transfected with 500 ng pHIV-1 or pHIV-1_EnvCpG86-561_ and 500 ng pGFP-FLAG. RNA was extracted from cell lysates **(A)** and media **(B)** and genomic RNA (gRNA) abundance was quantified by qRT-PCR. The bar charts show the average value of three independent experiments normalized to the value obtained for HeLa Control CRISPR cells co-transfected with pHIV-1 and pGFP-FLAG. Data are represented as mean ± SD. *p < 0.05 as determined by a two-tailed unpaired t-test. p-values for HIV-1_EnvCpG86-561_ genomic RNA in Control cell versus ZAP-G1, KHNYN-G1 Bulk, KHNYN-G1 Clone A, and KHNYN-G1 Clone B CRISPR cell lysates are 2.98 x 10^-2^, 3.70 x 10^-2^, 4.26 x 10^-2^, and 1.30 x 10^-4^, respectively. p-values for HIV-1_EnvCpG86-561_ genomic RNA in Control cell versus ZAP-G1, KHNYN-G1 Bulk, KHNYN-G1 Clone A, and KHNYN-G1 Clone B CRISPR cell media are 1.65 x 10^-2^, 8.64 x 10^-3^, 2.82 x 10^-3^, and 3.54 x 10^-4^, respectively.

Finally, we titrated CRISPR-resistant pKHNYN-1 or pKHNYN-2 into the KHNYN CRISPR cells. Even very low levels of KHNYN-1 or KHNYN-2 restored selective inhibition of HIV-1_EnvCpG86-561_ in these cells (Figure 9A-B) and KHNYN-1 was consistently slightly more active than KHNYN-2. This shows that both isoforms are capable of inhibiting infectious virus production of HIV-1 containing clustered CpG dinucleotides and also confirms the specificity of the KHNYN knockdown. We also analyzed whether KHNYN with the KH-like domain deleted or mutations in the NYN domain could inhibit HIV-1_EnvCpG86-561_ infectious virus production in the CRISPR cells. 31.25 ng of KHNYN-1 inhibited HIV-1_EnvCpG86-561_ to similar levels as in the Control CRISPR cells (Figure 9C-D) and all of the mutations substantially reduced KHNYN antiviral activity. Overall, endogenous KHNYN is required for CpG dinucleotides to inhibit HIV-1 infectious virus production.

**Figure 9.**
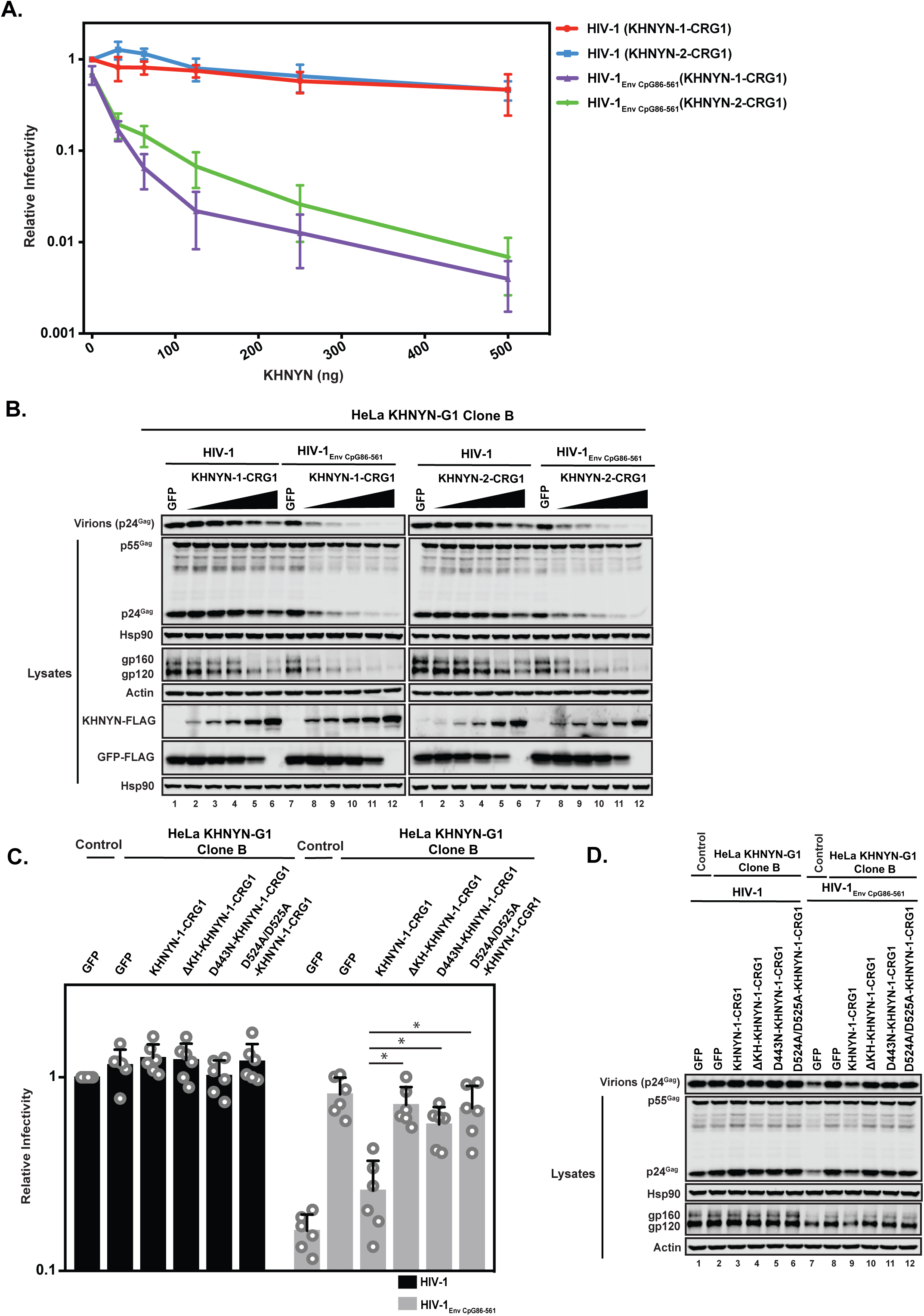
Deletion of the KH-like domain or mutations in the NYN domain in KHNYN prevent reconstitution of CpG-dependent inhibition of HIV-1 infectious virus production in KHNYN CRISPR cells. **(A-B)** HeLa KHNYN-G1 CRISPR clone B cells were transfected with 500 ng pHIV-1 or pHIV-1_EnvCpG86-561_ and 500 ng of pGFP-FLAG or 31.25 ng, 62.5 ng, 125 ng, 250 ng or 500 ng of pKHNYN-1-CRG1-FLAG or pKHNYN-2-CRG1-FLAG plus the amount of pGFP-FLAG required to make 500 ng total. Culture supernatants from the cells were used to infect TZM-bl reporter cells **(A)**. Each point shows the average value of three independent experiments normalized to the value obtained for HeLa Control CRISPR cells co-transfected with pHIV-1 and pGFP. Data are represented as mean ± SD. Gag expression in the media as well as Gag, Hsp90, Env, Actin, KHNYN-FLAG and GFP-FLAG expression in the cell lysates was detected using quantitative immunoblotting **(B)**. **(C-D)** HeLa KHNYN-G1 CRISPR clone B cells were transfected with 500 ng pHIV-1 or pHIV-1_EnvCpG86-561_, 468.75 ng of pGFP-FLAG and 31.25 ng of pKHNYN-1-CRG1-FLAG expressing either wild type or mutant proteins. Culture supernatants from the cells were used to infect TZM-bl reporter cells **(C)**. Each point shows the average value of six independent experiments normalized to the value obtained for HeLa Control CRISPR cells co-transfected with pHIV-1 and pGFP. Data are represented as mean ± SD. *p < 0.05 as determined by a two-tailed unpaired t-test. p-values for HIV-1_EnvCpG86-561_ with KHNYN-1-CRG1 versus ΔKH-KHNYN-1-CGR1, D443N-KHNYN-1-CGR1 and D524A/D525A-KHNYN-1-CGR1 are 2.40 x 10^-4^, 1.17 x 10^-3^, and 1.21 x 10^-3^. respectively. Gag expression in the media as well as Gag, Hsp90, Env, Actin, KHNYN-FLAG and GFP-FLAG expression in the cell lysates was detected using quantitative immunoblotting **(D)**.

## Discussion

Several members of the CCCH zinc finger domain protein family target viral and/or cellular mRNAs for degradation (Fu and Blackshear, 2017). For example, ZC3H12A degrades pro-inflammatory cytokine mRNAs and also inhibits viral replication of several different viruses, including HIV-1 and hepatitis C virus (Lin et al., 2013; Lin et al., 2014; Liu et al., 2013; Matsushita et al., 2009). It contains a CCCH zinc finger domain as well as a NYN endonuclease domain, which allows it to degrade specific RNAs (Matsushita et al., 2009; Xu et al., 2012). ZAP has four CCCH zinc finger domains and specifically interacts with CpG dinucleotides in RNA (Gao et al., 2002; Guo et al., 2004; Takata et al., 2017). However, it does not contain nuclease activity. While ZAP has been reported to directly or indirectly interact with components of the 5’-3’ and 3’-5’ degradation pathways including DCP1-DCP2, XRN1, PARN and the exosome, knockdown of several proteins in these pathways did not substantially rescue infectious virus production of HIV-1 containing clustered CpG dinucleotides (Goodier et al., 2015; Guo et al., 2007; Takata et al., 2017; Zhu et al., 2011). Therefore, we hypothesized that ZAP may interact with additional unidentified proteins that regulate viral RNA degradation.

Herein, we have identified that KHNYN is an essential ZAP cofactor that inhibits HIV-1 gene expression and infectious virus production when the viral RNA contains clustered CpG dinucleotides. KHNYN overexpression inhibits genomic RNA abundance, Gag expression, Env expression and infectious virus production for HIV-1 containing clustered CpG dinucleotides. This activity requires ZAP and TRIM25. Furthermore, depletion of KHNYN using CRISPR-Ca9 specifically increased RNA abundance and infectious virus production for HIV-1 containing clustered CpG dinucleotides. This indicates that KHNYN is essential for CpG dinucleotides to inhibit infectious virus production. We hypothesize that, after ZAP is activated by TRIM25, a complex containing ZAP and KHNYN binds the viral CpG containing RNA. ZAP and KHNYN could directly interact to form a heterodimer or there could be other factors mediating this interaction. The interaction between ZAP and KHNYN has been detected using several different assays including yeast-2-hybrid, co-immunoprecipitation, affinity purification–mass spectrometry (Huttlin et al., 2017) and BioID (Youn et al., 2018). If there is an unknown factor mediating this interaction, it would have to present in the yeast-2-hybrid assay. It remains unclear how TRIM25 regulates ZAP binding to viral RNA, but it is not required for ZAP and KHNYN to interact. Interestingly, TRIM25 co-immunoprecipitates with KHNYN and the ZAP antiviral complex may simultaneously consist of all three proteins. ZAP and TRIM25 are interferon-stimulated genes while KHNYN is not induced by interferon in human cells (Shaw et al., 2017). Whether KHNYN is regulated by type I interferons or viral infection in a different way, such as post-translational modification, is not known.

The zinc finger RNA binding domains in ZAP could determine the sequence specificity to target KHNYN to CpG regions in viral RNA. This would allow the endonuclease domain in KHNYN to cleave this RNA, thereby inhibiting viral RNA abundance. Conceptually, the ZAP-KHNYN complex could function similarly to ZC3H12A, but with the RNA binding and endonuclease domains divided between the two proteins. The NYN domain in KHNYN could cleave HIV-1 RNA containing CpG dinucleotides similar to how ZC3H12A cleaves a specific site in the 3’ UTR of the IL-6 mRNA (Matsushita et al., 2009). While we do not yet have evidence that the NYN domain in KHNYN is an active endonuclease domain, it is highly conserved with the NYN domain in ZC3H12A and is required for KHNYN activity. Strikingly, mutation of two conserved aspartic acid residues in the NYN domain predicted to coordinate a magnesium ion necessary for nucleophilic attack of the target RNA eliminated KHNYN antiviral activity. However, biochemical and structural studies will be necessary to determine the specific nature of the interaction between ZAP, KHNYN, TRIM25 and RNA and how these interactions promote viral RNA degradation.

An increasingly common theme for RNA decay is that endonucleic and exonucleic degradation pathways work together to fully degrade RNAs. For example, nonsense-mediated decay (NMD) targets mRNAs that do not efficiently terminate translation at the stop codon and uses up to four mechanisms to degrade these mRNAs: endonucleic cleavage, deadenylation, decapping and exonucleic degradation (Lykke-Andersen and Jensen, 2015). In this pathway, the endonuclease SMG6 interacts with the core regulatory protein UPF1 and cleaves mRNA near a premature termination codon. The 5’ and 3’ cleavage fragments are then degraded by the 5’-3’ exonuclease XRN1 and the 3’-5’ exonuclease exosome complex. The (CCR4)–NOT deadenylase complex and DCP1-DCP2 decapping complex are recruited by proteins in the NMD complex including UPF1, SMG5, SMG7 and PNRC2. Similarly, the 5’-3’ and 3’-5’ degradation pathway components previously shown to interact with ZAP could work in conjunction with KHNYN-mediated endonucleic decay (Goodier et al., 2015; Guo et al., 2007; Zhu et al., 2011). In this model, KHNYN would initiate cleavage of the viral RNA and the exonucleic pathways would then degrade the resulting RNA fragments. Identifying the full complement of ZAP-interacting factors and characterizing how these target viral RNAs for degradation will be an exciting area of future investigation.

## Materials and Methods

### Plasmids

All primers were ordered from Eurofins. Polymerase chain reactions (PCR) for cloning steps were performed with Phusion High Fidelity polymerase (New England Biolabs). KHNYN-2 (NM_001290256) was synthesized by GenScript. KHNYN-1 was cloned by amplifying the nucleotides 123-2157 from KHNYN-2 and sub-cloning the PCR product into the pcDNA3.1 (+) backbone using the HindIII site in the vector and SbfI site in the KHNYN open reading frame. KHNYN-1 ΔKH, D443N, D524A/D525A, -CRG1, -CRG2 and KHNYN-2 ΔKH, D484N, D565A/D566A, -CRG1, -CRG2 were generated via overlap extension PCR and subsequently sub-cloning the PCR product into the pcDNA3.1 (+) backbone as described above. pGFP-FLAG was cloned by amplifying GFP from pcDNA3.1-GFP(Swanson et al., 2010) and cloning it into pcDNA3.1. Diagnostic restriction enzyme digestion and DNA sequencing (Eurofins, Genewiz) was used to ensure the correct identity of modified sequences inserted into plasmids.

To generate HIV-1_EnvCpG86-561_, we synthesized a HIV-1_NL4-3_ EcoRI/StuI DNA fragment with synonymous mutations that inserted 36 CpG dinucleotides into *env* (Figure 2– figure supplement 1). This DNA fragment was digested with *EcoRI* and *StuI* and inserted into the corresponding sites of pHIV-1_NL4-3_ in pGL4 (Antzin-Anduetza et al., 2017).

### Cell lines

HeLa and HEK293T cells were obtained from the ATCC and were maintained in high glucose DMEM supplemented with GlutaMAX, 10% fetal bovine serum, 20µg/mL gentamicin or 100 U/ml penicillin and 100 μg/ml streptomycin and incubated with 5% CO_2_ at 37°C. CRISPR guides targeting the firefly luciferase gene, lacZ gene, human TRIM25, ZAP (also known as ZC3HAV1), and KHNYN genes were cloned into *BsmB*I restriction enzyme sites in the lentiviral vector genome plasmid lentiCRISPRv2(Sanjana et al., 2014). The CRISPR guide sequences are: LacZ-G1: 5’-CGA TTA AGT TGG GTA ACG CC-3’, Luciferase-G1: 5’-CTT TAC CGA CGC ACA TAT CG-3’, TRIM25-G1: 5’-GAG CCG GTC ACC ACT CCG TG-3’, ZAP-G1: 5’-ACT TCC ATC TGC CTT ACC GG-3’, KHNYN-G1: 5’-GGG GGT GAG CGT CCT TCC GA-3’, KHNYN-G2: 5’-CAG ACA CCG CAA AGC GAT CT-3’. LentiCRISPR vectors encoding guides RNA targeting KHNYN or LacZ were produced in HEK293T cells seeded in a 10cm dish and transfected using 100 ul PEI with 8µg of lentiCRISPRv2-Guide, 8µg of pCRV1-HIV-Gag Pol(Neil et al., 2008) and 4µg of pCMV-VSV-G(Neil et al., 2008). Lentiviral vectors encoding guide RNAs targeting ZAP or TRIM25 were produced by transfecting HEK293T cells seeded in a 6-well plate using 10 µl PEI with 0.5µg pVSV-G(Fouchier et al., 1997), 1.0 µg pCMVΔR8.91(Zufferey et al., 1997), and 1.0 µg LentiCRISPRv2-Guide. Virus containing supernatant was harvested 48-hr after transfection, rendered cell-free via filtration through 0.45 µM filters (Millipore) and used to transduce HeLa or HEK293T cells followed by selection in puromycin.

### Yeast two-hybrid screen

The yeast two-hybrid screen was performed by Hybrigenics Services (ULTImate Y2H, www.hybrigenics-services.com). Full length ZAP-S, ZAP-L and KHNYN-2 were used as LexA-bait fusion proteins (pB27; N-LexA-bait-C fusion) to screen the Induced Macrophages library. Because ZAP-L and ZAP-S were neither toxic nor autoactivating, a selective medium without 3-Aminotriazol was used for the final screen. KHNYN-2 had some autoactivating activity the final screen used a selection medium supplemented with 10 mM 3-Aminotriazol.

### Transfections

HeLa and HEK293T cells were grown to 70% confluence in six-well plates. HeLa cells were transfected according to the manufacturer’s instructions using *Trans*IT®-LT1 (Mirus) at the ratio of 3 µL TransIT®-LT1 to 1 µg DNA. HEK293T cells were transfected according to the manufacturer’s instructions using PEI (1mg/mL) (Sigma-Aldrich) at the ratio of 4 µL PEI to1 µg DNA. For each transfection, 0.5µg pHIV-1 and the designated amount of pKHNYN-FLAG, pGFP-FLAG or pGFP(Swanson et al., 2010) were used for a total of 1µg DNA. The transfection medium was replaced with fresh medium after a 6-hr incubation (HEK293T) or 24-hr incubation (HeLa).

### Analysis of protein expression by immunoblotting

HeLa cells were seeded on 6-well plates and transfected the following day with as described above. Approximately 48-hr post-transfection, HeLa cells were lysed in radioimmunoprecipitation assay (RIPA) buffer (150 mM NaCl, 1 mM EDTA, 1% Triton X-100, 1% sodium deoxycholate, 0.1% SDS, 10 mM Tris–HCl pH 7.5) supplemented with complete protease inhibitor (Roche) and sheared using a 0.5G needle. Media was filtered through a 0.45 µM filter and virions were pelleted for 2-hr at 20,000 x g through a 20% sucrose cushion in phosphate-buffered saline (PBS) solution. The pellet was resuspended in 2X loading buffer (60 mM Tris–HCl pH 6.8, 2% sodium dodecyl sulfate (SDS), 10% glycerol, 10% β-mercaptoethanol, 0.1% bromophenol blue). Cell lysates and virions were resolved by 10 % SDS-polyacrylamide gel electrophoresis (PAGE), transferred to a nitrocellulose membrane (GE Healthcare) and blocked in 1% non-fat milk in PBS with 0.1% Tween 20 (Fischer Bioreagents). Primary antibodies were incubated for 2 hours at room temperature. After washing in PBS, blots were incubated for 1 hour with the appropriate secondary antibody. Bound antibodies were visualized on the LI-COR^TM^ (Odyssey Fc) measuring the immunofluorescence or using Amersham ECL Prime Western Blotting Detection reagent (GE Lifesciences) for HRP-linked antibodies using an ImageQuant^TM^ (LAS4000 Mini). For the co-immunoprecipitation experiments, lysates were resolved using precast Mini-PROTEAN TGX gels 8-16% gradient gels (Bio-Rad) and transferred to nitrocellulose membranes (Bio-Rad). Antibodies used in study were 1:50 HIV-1 anti-p24Gag(Chesebro et al., 1992), 1:3000 anti-HIV-1 gp160/120 Rabbit (ADP421; Centralized Facility for AIDS Reagents (CFAR)), 1:10,0000 anti-HSP90 (sc7947, Santa Cruz Biotechnology), 1:5000 anti-HSP90 Rabbit (GeneTex, GTX109753), 1:4000 anti-HSP90 Mouse (SantaCruz, sc-515081), 1:5000 anti-ZAP (Abcam, ab154680), 1:1000 anti-β-Actin Mouse (Sigma, A2228), 1:2500 anti-DYKDDDDK (Rabbit) (Cell Signaling, 14793), 1:2500 anti-FLAG (Mouse) (Sigma, F1804), 1:2500 anti-FLAG (Rabbit) (Sigma, F7425), 1:10000 anti-TRIM25 (Abcam, ab167154), 1:10,000 Dylight™ 800-conjugated secondary antibodies (Cell Signaling Technology, 5151S and 5257S), 1:5000 anti-rabbit HRP (Cell Signaling Technology, 7074), 1:5000 anti-mouse HRP (Cell Signaling Technology, 7076), 1:4000 anti-rabbit IRDye 800CW (LI-COR, 926-32211) or 1:4000 anti-mouse IRDye 680RD (LI-COR, 926-68070).

### TZM-bl infectivity assay

Media was recovered approximately 48-hr post-transfection and cell-free virus stocks were generated by filtering the media through 0.45 µM filters (Millipore). The TZM-bl indicator cell line was used to quantify the amount of infectious virus(Derdeyn et al., 2000; Platt et al., 1998; Wei et al., 2002). TZM-bl cells were seeded at 70% confluency in 24-well plates and infected by overnight incubation with virus stocks. 48-hr post infection, the cells were lysed and infectivity was measured by analyzing β-galactosidase expression using the Galacto-Star™ System following manufacturer’s instructions (Applied Biosystems). β-galactosidase activity was quantified as relative light units per second using a PerkinElmner Luminometer.

### Immunoprecipitation assays

HEK293T cells in 6-well plates were transfected with 800ng of pKHNYN-1-FLAG, pKHNYN-2-FLAG, pGFP-FLAG as a control using 3 µL TransIT®-LT1 per 1µg of DNA added. For the experiments in which lysates were treated with Ribonuclease A (RNase A), 500ng of pHA-ZAP-L (Kerns et al., 2008) was also added. The cells were lysed on ice in lysis buffer (0.5% NP-40, 150mM KCl, 10mM HEPES pH 7.5, 3mM MgCl_2_) supplemented with complete EDTA-free Protease inhibitor cocktail tablets (Sigma-Aldrich), 10mM N-Ethylmaleimide (Sigma-Aldrich) and PhosSTOP tablets (Sigma-Aldrich). The lysates were sonicated and then centrifugated at 20,000 x g for 5 minutes at 4°C. 50μl of the post-nuclear supernatants was saved as the Input lysate and 450μl were incubated with either 18μg of anti-Flag antibody (Sigma-Aldrich, F7425) or 4.275μg of anti-ZAP antibody (Abcam) for one hour at 4°C with rotation. Protein G Dynabeads (Invitrogen) were then added and incubated for three hours at 4°C with rotation. The lysates were then washed 4 times with wash buffer (0.05% NP-40, 150mM KCl, 10mM HEPES pH 7.5, 3mM MgCl_2_) before bound proteins were eluted with Laemmli buffer and boiled for 10 minutes. When indicated, RNase A (Sigma-Aldrich) was added to the post-nuclear supernatant and incubated for 30 minutes at 37°C. Protein expression was analyzed via Western blot as described above.

### RNA Purification and quantitative RT-PCR

Total RNA was isolated from transfected HeLa cells using a QIAGEN RNAeasy kit accordingly with the manufacture instructions. Viral RNA was extracted from cell supernatants using a QIAGEN Qiamp Viral mini kit accordingly with the manufacture instructions. 500 ng of purified cellular RNA was reverse transcribed using random hexamer primers and a High Capacity cDNA Reverse Transcription kit (Applied Biosystems). Quantitative PCR was performed using a QuantiStudio 5 System (Thermo Fischer). Relative quantification was determined using the GAPDH Taqman Assay (Applied Biosystems, Cat# Hs99999905_m1) and absolute quantification was determined using a standard curve of the HIV-1 provirus DNA plasmid. The genomic RNA primers were GGCCAGGGAATTTTCTTCAGA / TTGTCTCTTCCCCAAACCTGA (forward/reverse) and the probe was FAM-ACCAGAGCCAACAGCCCCACCAGA-TAMRA.

### Microscopy

Cells were seeded on 24-well plates on coverslips pre-treated with poly-lysine. HEK293T cells expressing a control guide RNA targeting the LacZ gene or a guide RNA targeting ZAP were transfected with 250ng of pKHNYN-FLAG. 24 hours post-transfection, the cells were fixed with 4% paraformaldehyde for 15 minutes at room temperature, washed with PBS, and then washed with 10mM glycine. The cells were then permeabilized for 15 minutes with 1% BSA and 0.1% Triton-X in PBS. Mouse anti-FLAG (1:500) and rabbit anti-ZAP (1:500) antibodies were diluted in PBS/0.01% Triton-X and the cells were stained for one hour at room temperature. The cells were then washed 3 times in PBS/0.01% Triton-X and incubated with Alexa Fluor 594 anti-mouse and Alexa Fluor 488 anti-rabbit antibodies (Molecular Probes, 1:500 in PBS/0.01% Triton-X) for 45 minutes in the dark. Finally, the coverslips were washed three times with PBS/0.01% Triton-X and then mounted on slides with Prolong Diamond Antifade Mountant with DAPI (Invitrogen). Imaging was performed on a Nikon Eclipse Ti Inverted Microscope, equipped with a Yokogawa CSU/X1-spinning disk unit, under 60-100x objectives and laser wavelengths of 405nm, 488nm and 561nm. Image processing and co-localization analysis was performed with NIS Elements Viewer and Image J (Fiji) software.

### Statistical analysis

Statistical significance was determined using unpaired two-tailed t tests calculated using Microsoft excel software. Data are represented as mean ± SD. Significance was ascribed to p values p < 0.05.

## ACKNOWLEDGEMENTS

We thank other members of the Neil and Swanson laboratories for helpful discussions. The following reagents were obtained through the NIH AIDS Research and Reference Reagent Program, Division of AIDS, NIAID, NIH: TZM-bl from Dr. John C. Kappes, Dr. Xiaoyun Wu and Tranzyme Inc; HIV-1 p24 Hybridoma (183-H12-5C) from Dr. Bruce Chesebro. The Antiserum to HIV-1 gp120 #20 (ARP421) was obtained from the NIBSC Centre for AIDS Reagents. Dr Harmit Malik kindly provided the ZAP-L expression vector. These studies were funded by MRC Discovery Award MC/PC/15068 and a Wellcome Trust Senior Research Fellowship (WT098049AIA) to SJDN and Medical Research Council grant MR/M019756/1 to CMS. MF is supported by the UK Medical Research Council (MR/R50225X/1) and is a King’s College London member of the MRC Doctoral Training Partnership in Biomedical Sciences. This work was also supported by the Department of Health via a National Institute for Health Research Comprehensive Biomedical Research Centre award to Guy’s and St. Thomas’ NHS Foundation Trust in partnership with King’s College London and King’s College Hospital NHS Foundation Trust.

## SUPPLEMENTAL FIGURE LEGENDS

**Figure 2–figure supplement 1.**
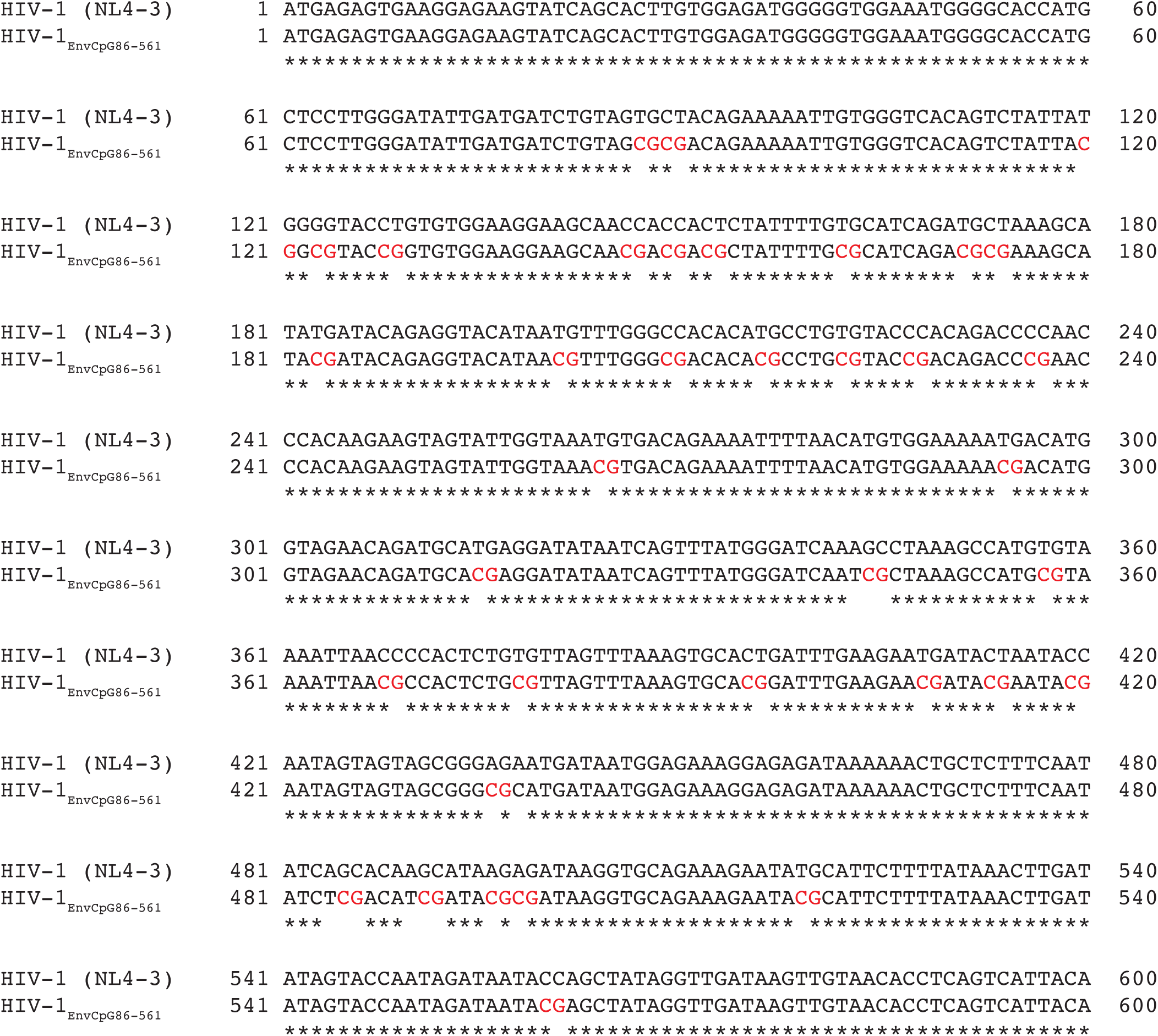
HIV-1_EnvCpG86-561_ contains 36 introduced CpG dinucleotides. MacVector ClustalW alignment of nucleotides 1-600 of *env* from wild type HIV-1 and HIV-1_EnvCpG86-561_. CpG dinucleotides introduced through synonymous mutations are highlighted in red.

**Figure 6–figure supplement 1.**
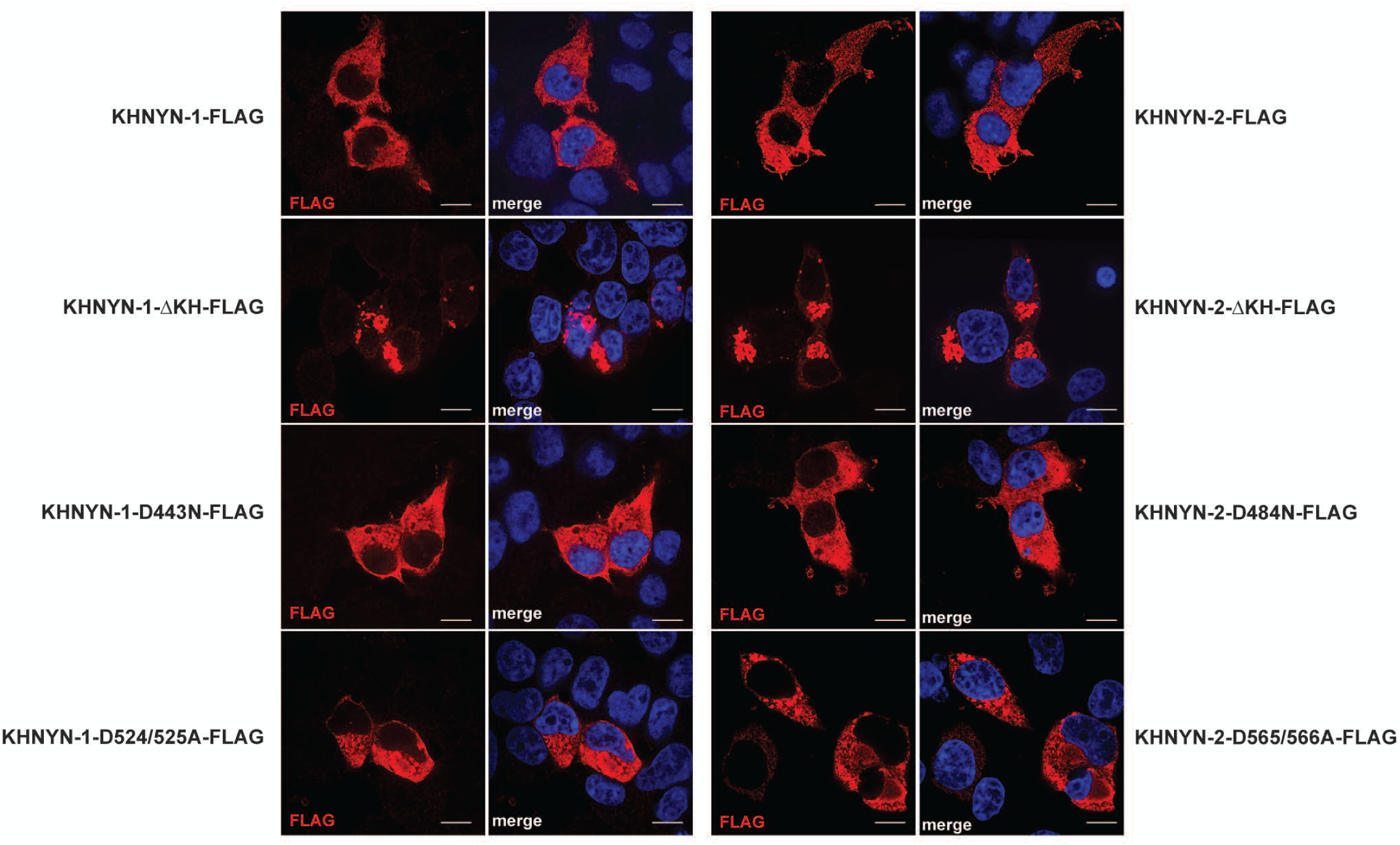
KHNYN mutant proteins localize to the cytoplasm. Panels show representative fields for the localization of KHNYN-1/2-FLAG wild type or mutant proteins in 293T Control CRISPR cells. Cells were stained with an anti-FLAG antibody (red) and DAPI (blue). The scale bar represents 10 µM.

**Figure 6–figure supplement 2.**
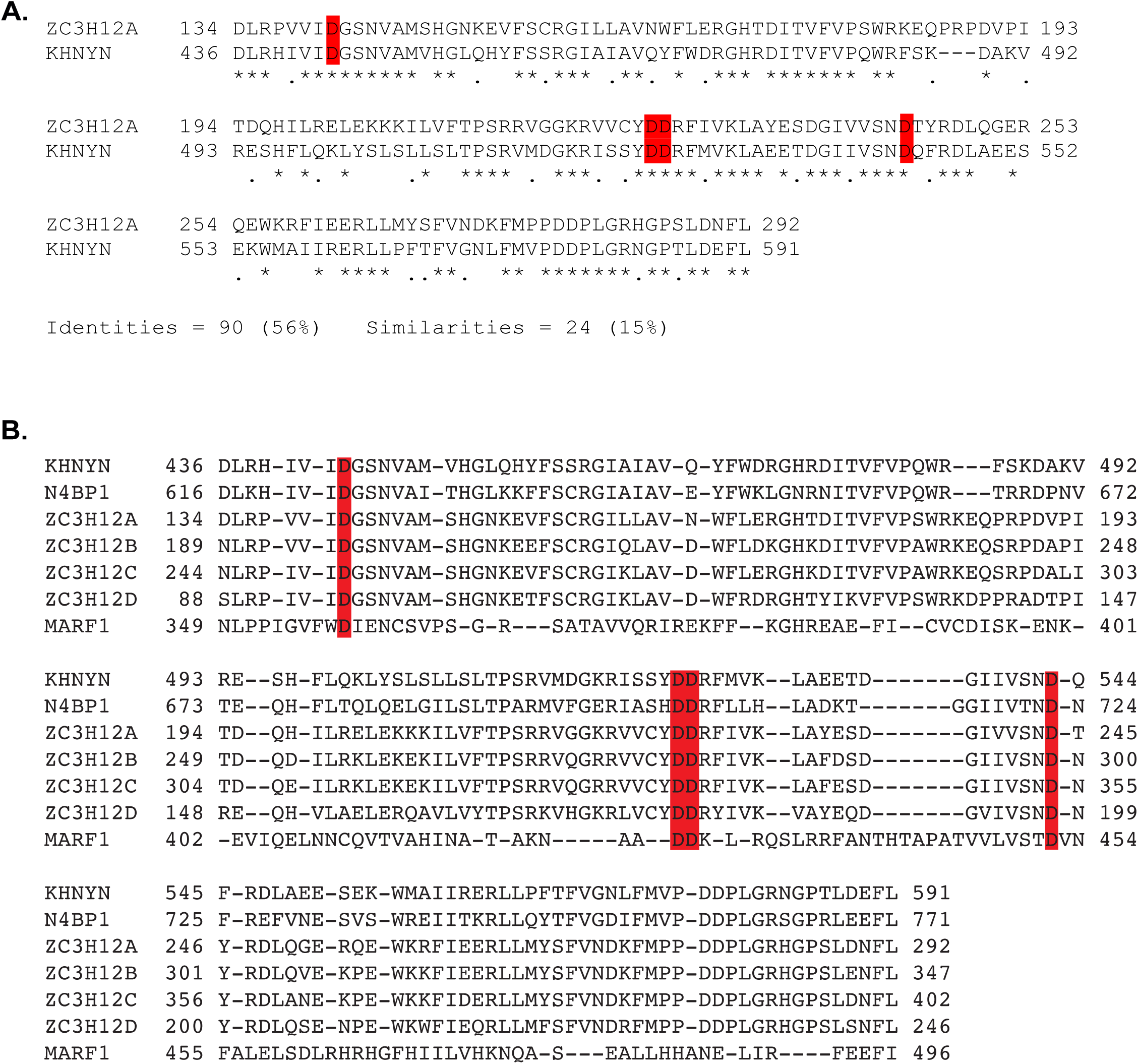
NYN domain alignment. **(A)** MacVector ClustalW alignment of the NYN domains from KHNYN and ZC3H12A. The amino acid identity and similarity is shown. The four aspartic acid residues that coordinate the magnesium ion in the active site are highlighted in red. **(B)** MacVector ClustalW alignment of NYN domains from KHNYN, N4BP1, ZC3H12A, ZC3H12B, ZC3H12C, ZC3H12D and MARF1. The four aspartic acid residues that coordinate the magnesium ion in the active site are highlighted in red.

